# Six decades of change across all North American bird interaction networks

**DOI:** 10.64898/2026.04.30.721628

**Authors:** Phoebe L. Zarnetske, Stéphane Dray, Vincent Miele

## Abstract

Species interactions are essential for maintaining biodiversity in the face of global changes. Yet lack of interaction data has hindered progress in understanding spatiotemporal changes in ecological network structure and function. To address this shortfall, we perform network analysis on a continental-scale metanetwork of bird interactions (AvianMetaNetwork), and quantify shifts in trophic, commensalism, parasitism, competition, mobbing, and facilitation networks using 13762 interactions among 687 North American birds. Since 1970, changes in species’ abundance and human activity drove network shifts, especially in eastern North America. With this metanetwork, we uncovered more than a half century of previously undocumented shifts in the structure of each avian network at a continental scale, with large implications for the functioning of bird communities and the ecosystems that rely on them.

## Main Text

Every species interacts with other species, directly, and indirectly through different network relationships involving multiple types of positive, negative, and neutral interactions (*1*, *2*). Species interactions are fundamental drivers of species’ distributions (*1*, *3*) and can affect species’ occurrence at multiple spatial scales (*4–6*). These interactions reflect eco-evolutionary processes underlying the structure and function of ecological communities and are vital to maintaining ecological community integrity in the face of global changes (*7*). Species interactions are also critical to informing species’ threat status and reserve design (*8*, *9*), and reaching conservation and management goals (*10*, *11*), including achieving 30×30 Global Biodiversity Framework targets (*12*). However, global changes are reshuffling, enhancing, and diminishing species interactions and the networks they form (*13–17*), thus affecting species distributions. It is therefore critical to quantify how species interactions and their networks are changing through space and time (*7*, *18–21*) to predict the future of biodiversity.

However, a lack of comprehensive interaction data has hindered progress (*21–23*). Indeed, it is practically impossible to collect comprehensive field-based observations of interactions at broad spatial and temporal scales. To address this challenge, researchers create metanetworks (*24*), which are networks of potential interactions that incorporate knowledge of empirically observed or ecologically feasible interactions among the species present in a pre-defined region (*25*). The concept of the metanetwork can be viewed as a biotic equivalent of the fundamental niche (*26*, *27*) representing the maximum potential interaction space before other filters (e.g., environment) act on local communities. Filtering these metanetworks by spatio-temporal community data such as presence/absence (*28*) or abundance data (*29*), allows us to generate realized local networks, thus providing a unique opportunity to track changes in network structure and function across space and time.

Birds are ideal taxa to address knowledge gaps in regional to continental-scale ecological networks (*30–33*) as they participate in a wide range of interactions (*34–36*) that have been extensively documented over the years. By synthesizing text-based natural history (*33*, *37*) into metanetworks, researchers can go beyond species level analysis to uncover community and network-level dynamics. In these metanetworks, species may occupy redundant interaction niches characterized by similar interactions with similar partners (*38*, *39*) providing robustness of functions against species turnover. This redundancy, exemplified for birds by host-sharing in brood parasites (*40*), nest webs involving primary and secondary cavity nesting species (*41*), or collective foraging in mixed-species flocks (*36*, *42*), ensures that key functions persist in communities even if individual species decline. Hence, integrating functional redundancy in spatial or temporal trends analysis may reveal shifts in functions that cannot be detected by species-level analysis (*29*).

Here, we leverage the AvianMetaNetwork (*37*, *43*), a comprehensive empirically-based database of six interaction types (trophic, commensalism, parasitism, competition, mobbing and facilitation), to quantify 60 years of bird community restructuring across 32 North American ecoregions. With these data, we build on prior studies of avian biodiversity at continental scales (e.g., (*44*, *45*)) and species-level population changes over time (e.g., (*46–49*)) to improve understanding at the network level. First, using the North American Breeding Bird Survey bird count data, we quantify abundance turnover at the species level. Then, by applying network clustering (*50*) to avian metanetworks, we identify functional groups of species within each ecological network to quantify functional redundancy. By merging the abundance and network-based information, we quantify the network structure turnover across ecoregions and multiple decades. To understand the potential causes of turnover at the species-level and network-level, we quantified climate and human pressure trajectories, which may affect species’ abundance and network turnover given the documented effects of climate change (e.g.,(*51*, *52*)) and human impacts (e.g.,(*46*, *49*, *53*)) on individual species and avian diversity. Investigating these changes over these longer timeframes and broader spatial scales enabled us to uncover a wide range of potential responses to change, and identify networks and regions that are more vulnerable due to increased anthropogenic stressors. Ultimately, this knowledge can help inform prioritization efforts for species, ecological communities, and regions most at risk of losing important ecological roles and functions provided by birds.

## Results

### Continuous avian species abundance turnover since 1970 across North America

We found a large temporal shift from 1970 to 2023 in terms of avian community composition at the North American scale. Using multiple factor analysis (MFA) on the series (per ecoregion) of year-by-species abundance tables, we found that the first MFA axis explaining 29% of the species turnover corresponds to the timeline of successive years from 1970 through 2023 (Fig. 1A). Importantly, this trend emerged solely as a result of the MFA analysis, as time was not specified in the analysis itself. Although all ecoregions exhibit species’ abundance turnover, the link values (which measure the agreement of each ecoregion with the continental trend) capture the spatial variations in this turnover. Higher values are observed in the East than in the West (Fig. 1B), suggesting that the temporal shift observed at the continental scale is more pronounced in the eastern regions. The lower link values are observed in ecoregions in the far North and South. The pattern of the species normalized abundance confirms these variations (Fig. 1C,D) with a clear abundance shift over the years in ecoregions with higher link values.

**Fig. 1.**
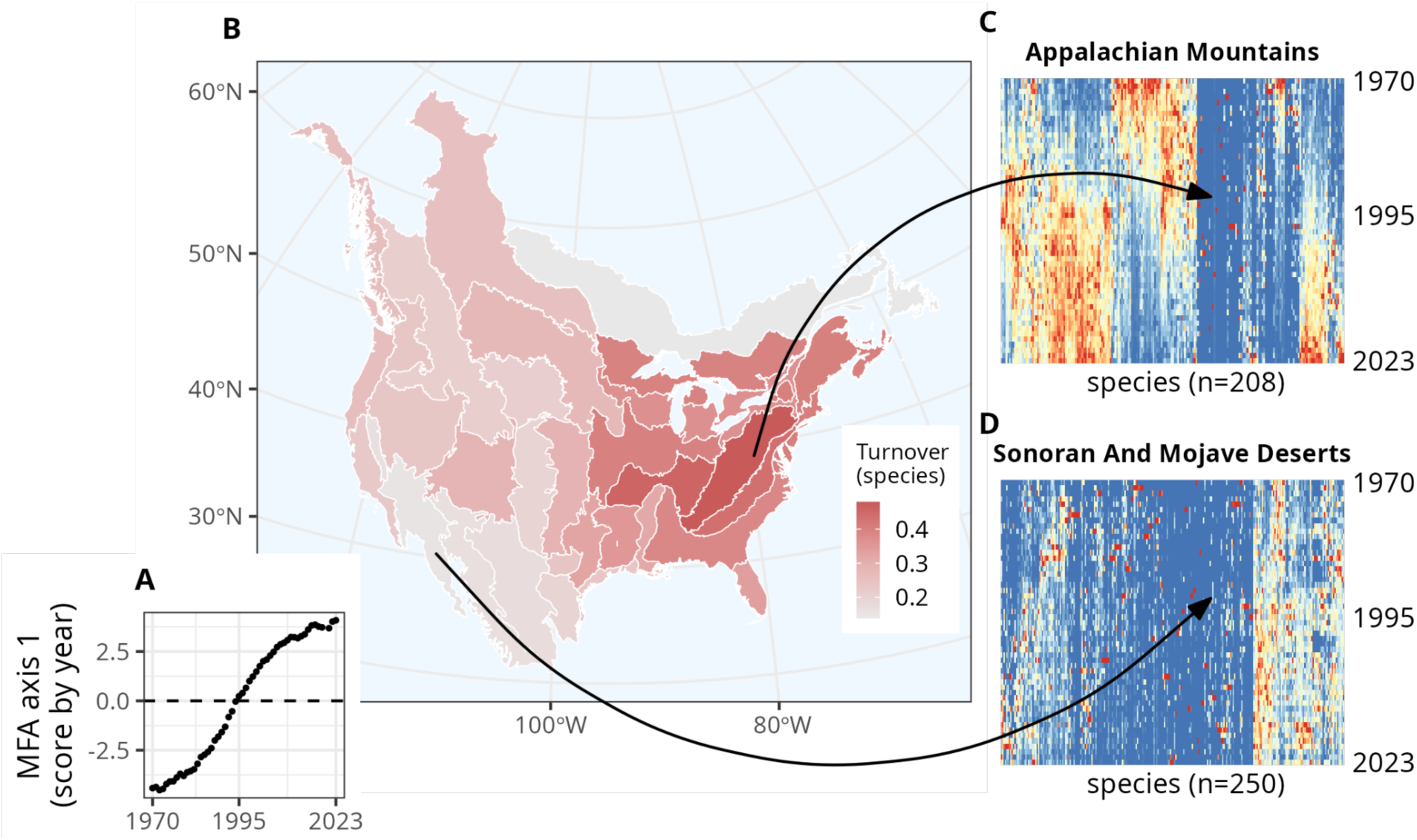
Species’ abundance turnover, by ecoregion. (**A**) Species-level MFA first axis showing continuous species’ relative abundance turnover over the entire time period, for the whole region (North America). (**B**) Map of the species’ relative abundance turnover by ecoregion, as expressed by the so-called link with the global Multiple Factor Analysis (MFA) first axis. (**C**) Heatmap showing species (columns) relative abundance (high values in red, medium in yellow, low in blue) for each subsequent year (rows) for the Appalachian Mountains ecoregion, which has a high link value. (**D**) Same as (**C**) for the Sonoran and Mojave Deserts ecoregion, which has a low link value. (**C-D**) The number of columns and the column ordering are different for the two figures; the column ordering was computed using a hierarchical clustering.

Species did not contribute to the temporal trend in community composition in the same way. We computed the score of each species on the first MFA axis and found that a subset of species have higher negative or positive score values (Fig. 2E). These species have a larger increase (Fig. 2C,F for example) or decrease (Fig. 2A,D for example) in abundance over time. These effects were more pronounced in ecoregions where the link to the continental trend was high (darker curves in Fig. 2), indicating that compositional shifts are driven by the species with higher variations in abundance over time.

**Fig. 2.**
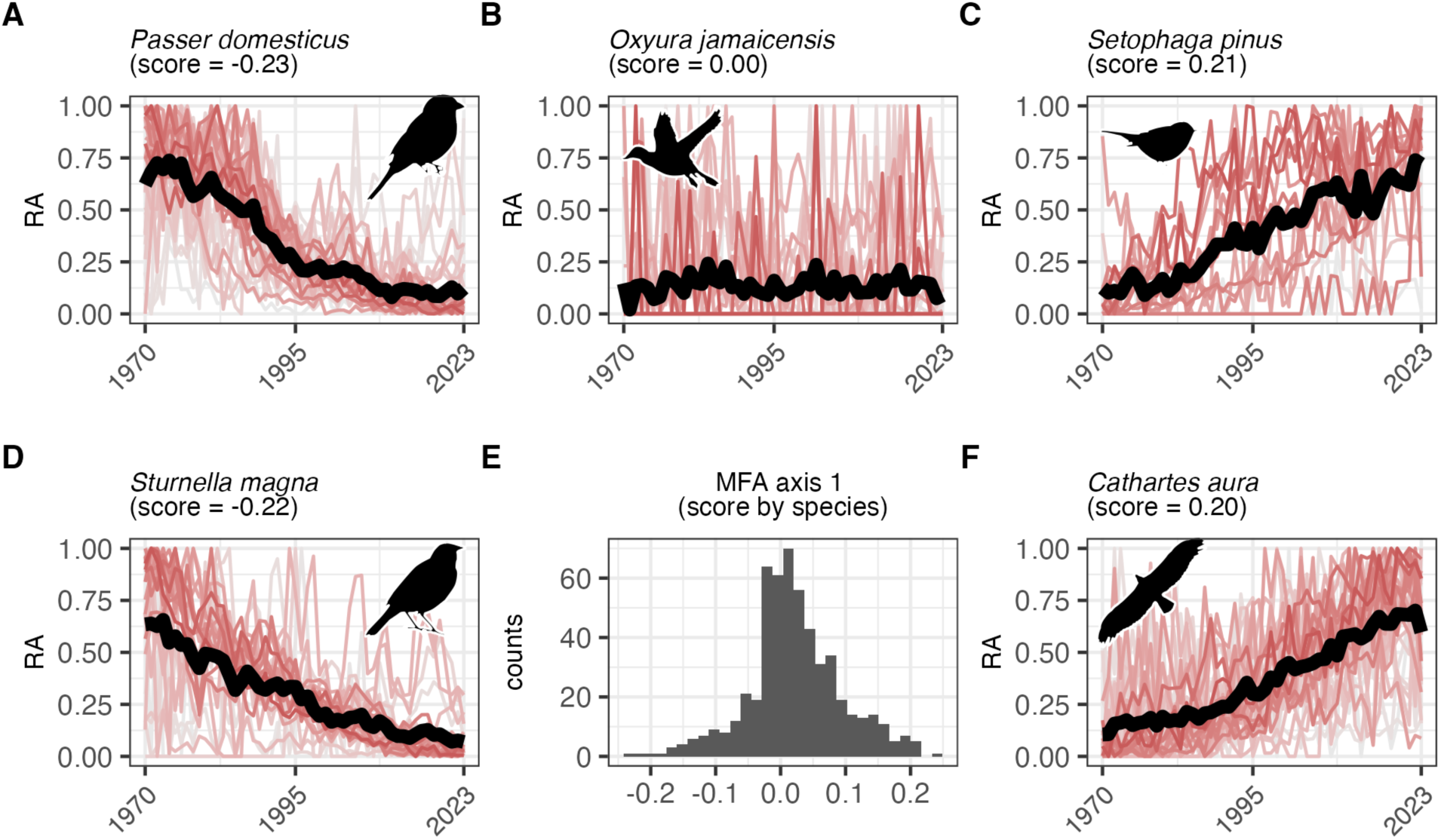
Example species’ relative abundance (RA) trends from 1970-2023 and MFA score. All the curves (in **A**,**B**,**C**,**D**,**F**) represent the different BCR ecoregions, with a color shade given by the turnover (link) value from Fig. 1 and the black and bold line corresponds to the average trend for the whole study area. Species with the highest negative MFA first axis scores (**A,D**) are more present earlier in the time period and species with the highest positive scores (**C,F**) are more present later in the time period. If the MFA analysis first axis score of a species is close to 0 (**B**), its relative abundance does not vary much over time, whereas if a species has a positive or negative score, it has more abundance turnover over time. Histogram (**E**) of the MFA first axis score for all species, showing the distribution of each species’ relative abundance trend over the entire time period, for the whole region (North America). Species’ common names are as follows: (**A**) *Passer domesticus* (House Sparrow), (**B**) *Oxyura jamaicensis* (Ruddy Duck), (**C**) *Setophaga pinus* (Pine Warbler), (**D**) *Sturnella magna* (Eastern Meadowlark), and (**F**) *Cathartes aura* (Turkey Vulture). Icons are CC0 1.0 from phylopic.org.

### Uncovering network-based functional redundancy: When species share interaction niches

Most networks contained a majority of the 687 species (Fig. 3), indicating that many North American species provide multiple functions within their communities. Using network machine learning (Daudin et al. 2008) we deciphered three to four functional groups of species within each of the six networks (Fig. 3, colors, tables S1-S6); each group includes species with the same role in terms of interaction with the others sharing a common interaction niche or “network function”.

**Fig. 3.**
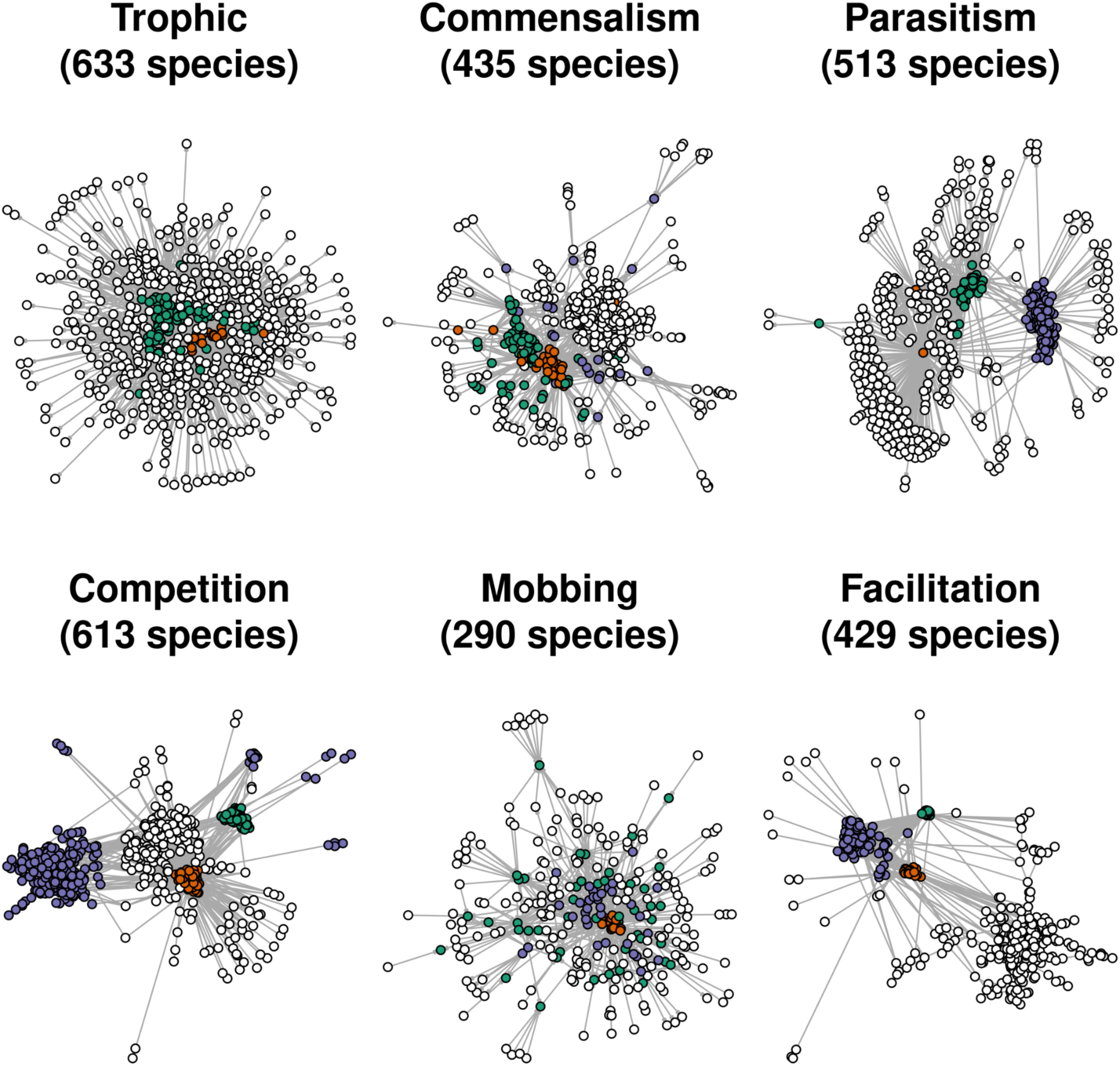
Network structure for the six types of interactions. Network representation showing the network groups (colors) obtained with stochastic block models, for all interacting species. Interaction directionality (small gray arrows) is shown except for the symmetrical networks (competition and facilitation). Colors and node positions are independent for the six networks, and node positions were obtained with the Fruchterman-Reingold algorithm. Species in white rarely interact with other species. Refer to tables S1-S6 for the probabilities of interaction between groups in each interaction type, and for species’ membership by interaction type and group. The groups retrieved by machine learning categorize trophic groups as top predators (orange), mesopredators (green), and prey (white); commensalism groups as providers (orange), beneficiaries (green, white, purple); parasitism groups as parasites (orange) and their different targets (white, green) and reciprocal parasites (purple); competition groups as reciprocal competitors (orange and green; purple to a lesser extent); mobbing groups as active mobbers (purple) and their predator targets (orange and green); and facilitation groups as group foragers (green), communal nesting and foraging (orange), or mixed flocking (purple).

The trophic network consists of 3 groups (colors in Fig. 3) that can be interpreted as top predators (orange), mesopredators (green) and prey (white). The commensalism network is organized around a first group (green) of species that use abandoned nests of other birds (including secondary cavity nesting birds), and receive most of these benefits from a group (orange) consisting of species that create nests (including primary cavity nesting species) or provide the majority of benefits to the other groups; other beneficiary groups include species that react to alarm calls of conspecifics or use their nests (white), or include species that mimic calls of other species (purple). The parasitism network is structured around a group of two hyperactive and generalist brood parasites (orange); *Molothrus ater* (Brown-headed Cowbird) and *Molothrus aeneus* (Bronzed Cowbird), whose target species’ nests are mostly in two other groups (white and green). Interestingly, there is another group of species (purple) where some species tend to be parasites of others in the same group (often through kleptoparasitism, e.g., gulls, raptors, corvids). Competition is dominated by modules of species in reciprocal competition over food, territory, and nesting sites (orange and green, and to a lesser extent, purple, with white rarely interacting with other groups). For mobbing, the network is dominated by a group of active mobbers (purple: predominantly passerines and colonial waterbirds) targeting two groups of predators (orange and green), one (orange: mostly apex species and scavengers) being composed of species targeting species of the same group. Finally, the facilitation network reveals a modular structure with 3 groups (all colors except white) being composed of species mainly facilitating each other through, for example, group foraging (green: mostly high latitude shorebirds and waterfowl), communal nesting and foraging (orange: mostly wetland and waterbird species like herons, egrets, ibis, spoonbills, storks, pelicans, cormorants), and mixed flocking (purple: mostly coastal seabirds, marine alcids, and shorebirds).

### Continuous shifts in network organization over time

The definition of groups of redundant species representing network functions is particularly informative because it goes beyond individual species turnover or abundance trends. We thus integrated these groups into the MFA analysis and obtained a similar temporal trend, with a first axis (39% of the total structure) containing years organized in a successive manner. This temporal trend was not imposed by the MFA method, but rather emerged as a result of the analysis (Fig. 4A). Similar to the species level case, we concluded that the group’s composition, i.e., the organization of the six networks, clearly shifted with time. However, whereas the turnover values were high for ecoregions in the East, the turnover values were lower in the West (Fig. 4B). As a result, the network organization tended to be more conserved in the Western regions (Fig. 4C,D).

**Fig. 4.**
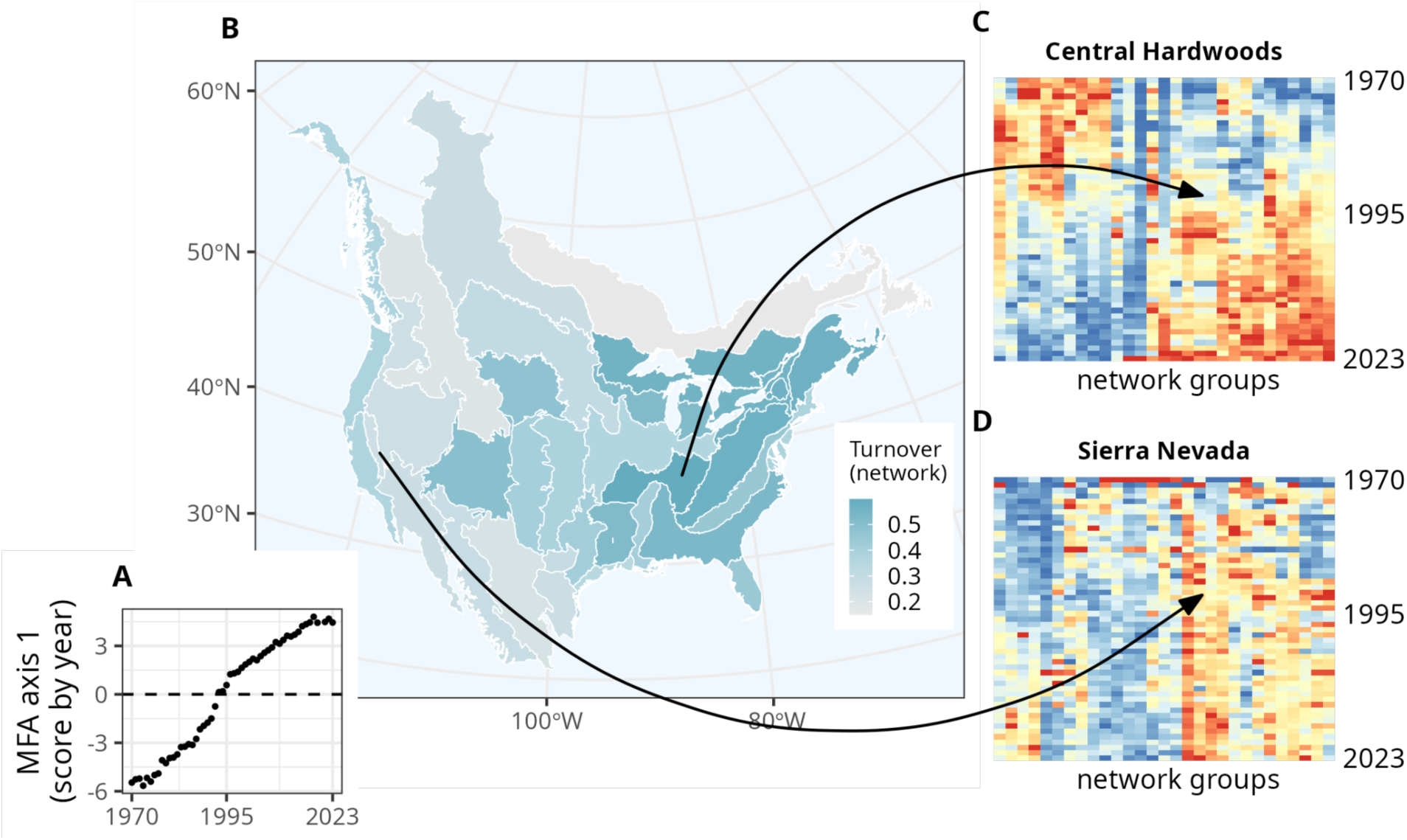
Network turnover, by ecoregion. (**A**) Network-level MFA first axis showing continuous network group relative abundance turnover over the entire time period, for the whole region (North America). (**B**) Map of the network group abundance turnover relative strength by BCR ecoregion, as expressed by the so-called link with the global Multiple Factor Analysis (MFA) first axis. (**C**) Heatmap showing groups (columns) abundance (high values in red, medium in yellow, low in blue) for each subsequent year (rows) for the Central Hardwoods ecoregion, which has a high link value. (**D**) Same as (**C**) for the Sierra Nevada ecoregion, which has a low link value. (**C-D**) The number of columns is constant but the column ordering is different for the two figures, and the column ordering was computed using a hierarchical clustering.

To investigate the amount of change per interaction type, we computed each interaction type’s contribution to the entire network-level turnover (i.e., changes in abundances between functional groups, fig. S1). This turnover was ubiquitous across all types of interaction network layers, all exhibiting similarly high levels of turnover. Since most species are involved in more than one type of interaction, changes in their abundance naturally spread across different networks.

Finally, we simultaneously mapped the turnover intensities since 1970 at the species and network levels (Fig. 5A). Both turnover intensities are consistent in most of the ecoregions (ascending diagonal colors of the Fig. 5A legend). Ecoregions experiencing the highest shifts in both species composition and network organization (i.e., higher link values) are primarily located in the East. However there are other configurations. In some regions, species abundance turnover does not translate as much into turnover at the network level (Fig. 5A red colors, e.g., New England/MidAtlantic Coasts and Eastern Tallgrass Prairie), indicating that network structure is more stable than the species community in these regions. In contrast, the opposite is observed, notably in Central regions (Fig. 5A blue colors, especially Badlands and Prairies) where moderate changes in species communities induced large network changes over this time.

**Fig. 5.**
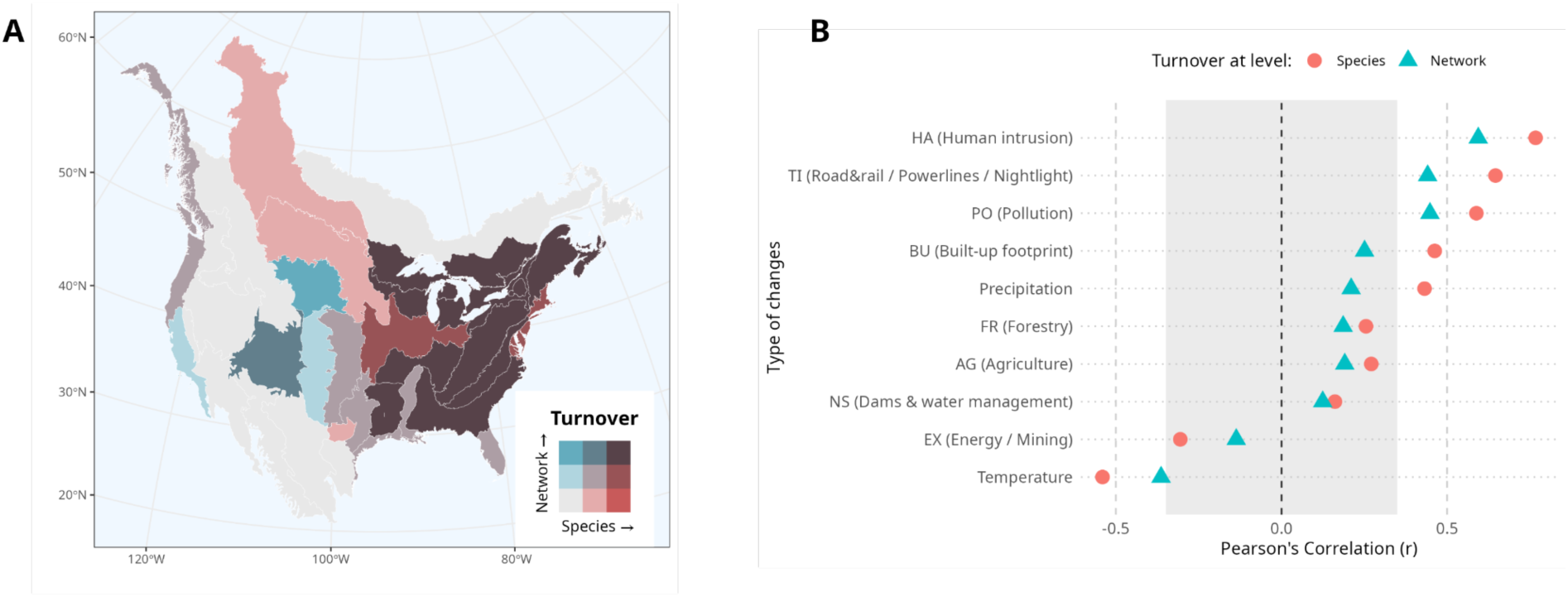
Species’ abundance versus network turnover, by ecoregion (A), and correlation between turnover at species or network level and climatic changes or human modification (B). (**A**) Bivariate mapping of species-based relative abundance turnover (link values from the species-level MFA; Fig. 1B) versus network-based turnover (link values from the network-level MFA; Fig. 4B). The color scheme is based on a RGB coding of the two turnover values, and higher turnover is in darker tones while lower turnover is in lighter tones. BCRs reflect all colors, except for the brightest red (highest species turnover and lowest network turnover). (**B**) Pearson correlation between species abundance and network turnovers with changes in climate and human modification (the gray area indicates non-significant values). Climatic changes (temperature and precipitation) were computed in the 1970-2023 period from the ERA5 re-analysis (*66*, *67*), whereas human modifications were restricted to the 1990-2020 period given the available timeframe of the Human Modification data (*58*, *68*).

### Human activities as the main drivers of abundance and network turnover

We observed a high Pearson correlation between the intensity of community changes (as measured by the MFA link values) and some of the human modification indexes (Fig. 5B, fig. S2). At the species and the network levels, we found that changes regarding human intrusion (HA), infrastructures (powerlines and nighttime lights, TI) and pollution (PA) are the most correlated with bird community shifts (correlations >0.50). In particular, the higher the HA modification, the higher species and network community shift. On the other hand, correlations with changes in climate are much lower and even negative for temperature changes. The latter observation suggests that regions with relatively less temperature change since 1970 (e.g., Peninsular Florida and Eastern Tallgrass Prairie) correspond to areas with large shifts in bird communities. Overall, these results suggest that human presence and footprint are likely the main drivers of turnover in bird species and networks.

## Discussion

By uncovering how avian network structures and functions change across nearly 6 decades at a continental scale, our study enhances prior knowledge about how and why North American avian populations shift over time and space (*45*, *46*, *48*, *49*, *52*, *53*). We leveraged a comprehensive database of empirically-based bird interactions in North America (*37*, *43*) and decades of bird observations (*54*) to discover clear temporal shifts in avian communities both at the species and network levels. Despite exceptions in some ecoregions, turnover in species’ abundance and networks are generally positively correlated across the continent, suggesting that changes in species composition induce changes in network functions. This reveals low functional redundancy in avian metanetworks where the decline in abundance of one species is not compensated by the increase of another species fulfilling the same ecological role. Hence, across nearly all ecoregions of North America, avian metanetworks are not robust to species turnover and are thus vulnerable to species decrease or loss.

All interaction networks are affected by these changes but the networks with the highest turnover are mobbing and commensalism (fig. S1). Spatially, the shifts are higher in the eastern United States (Fig. 4 darker shades) and some central or western ecoregions (Fig. 4 dark blue). Whereas traditional conservation policies are based on the presence and abundance of species of concern, our approach provides avenues to integrate interaction networks and ecological functions as conservation targets (as suggested by (*10*, *11*)). Indeed, we show that it is possible to evaluate functional redundancy using metanetworks, resulting in useful information to assist conservation prioritizations. For instance, ecoregions with low species turnover but high network turnover highlight areas where minor changes in species composition induce major changes in functions (Fig. 5A blue shades). This is especially the case with the Badlands and Prairies ecoregion which supports numerous breeding birds and species of conservation concern (*55*), and is more vulnerable due to its high network-level turnover (Fig. 5A). Thus, by using metanetworks to quantify functional redundancy, conservation objectives can shift from primarily focusing on protecting individual species populations to also protecting ecological networks, their functions, and the habitats that foster interactions.

Species-level turnover inevitably scales up to affect community turnover at the group level for several networks as most species are involved in more than one type of interaction so that shifts in their abundance naturally spread across different networks. Changes are strongest for mobbing interactions, which involve mainly predators and their potential prey that form the backbone of the organization of bird communities (*30*, *56*). As a telling example, we observed an increase in abundance of predators (see fig. S3), which potentially reflects the positive impact from conservation policies targeting birds of prey (e.g., raptors recovering after the DDT was banned; (*46*, *53*)). Targeting top predators is a success in a species-centric view but induces ubiquitous turnover across all types of interaction layers (e.g., trophic, mobbing, parasitism, competition). Consequently, isolated conservation measures targeting a single species or a single interaction type (such as a food web) can trigger unexpected cascading effects across network layers and fundamentally modify community organization.

The largest shifts in composition and associated function occurred in eastern North American ecoregions where human impacts have increased markedly over the last three decades. Although our analysis does capture changes in geographic distribution (as in (*45*) where latitudinal shifts, especially in the East, are documented with 209 individual ranges of North American birds), our results for 687 species provide new evidence of species-level and network turnover underlying large compositional changes.

Previous studies have focused on North American bird abundance change, with respect to changes in land cover and climate over 25-47 years (e.g., (*46*, *49*, *53*, *57*)). We build on these studies to enhance understanding of avian biodiversity change by providing the first continental assessment of avian network structure that maps directly to community functions over nearly six decades, and connecting these changes with climate and human modification impacts that align directly with biodiversity threats. For example, in 15 BCRs from 1990–2016, local bird community turnover in abundance at a scale of 1-5 BBS routes was likely driven by changes in land cover, whereas at the ecoregion scale, temperature rise explained more regional turnover (*57*). In another example, hotspots of bird abundance decline and faster rates in these declines for individual species abundances from 1987-2021 were likely driven by high-intensity agriculture across multiple U.S. regions (*49*). We investigated not only human activity footprint—which reflects land cover changes—but also human activity per se, using human modification change variables that map directly to IUCN threat status (*58*). We found that increased human intrusion (e.g., roads, railways, power lines, nighttime lights), human activity (e.g., use of roads, trails, and risk of invasive species), and pollution were more highly correlated with changes in bird species abundance and networks than the impact of other threats such as agricultural intensification or land cover change (as in (*49*, *57*)). Our findings identify a long-term signal of human activity on avian community structure and function, contributing evidence for the impacts of human activity and presence on wildlife, which has recently been highlighted as a concern, especially in less human-modified habitat (*59*, *60*). Although the high correlation with human activity does not equate to causality (*61*), prior studies point to the negative impact of human activity on bird survival and population size (e.g., (*46*, *53*)) which are demographic responses that underlie species abundance and network turnover.

Our metanetwork approach considers 13,762 potential interactions based on avian interactions that were directly observed and catalogued from a manual systematic review of literature (*43*). These interaction data made it possible to address questions at the continental scale and across a large taxonomic group. However, this approach includes several assumptions. First, the interactions were not directly observed among species through the BBS observations, as is the case for field studies of plant-pollinator networks (for example (*62*). In addition, these potential interactions were considered to be constant, despite the plasticity of interactions including the potential for reorganization, known as ‘rewiring’ (*63*). Also, groups were defined using the metanetwork and may not always reflect the actual role of species in the field. However, our approach considers potential interactions and is thus similar in reasoning to studies using the fundamental niche concept, which is used to describe a species’ ability to adapt to its environment (*26*).

We conducted our analysis at the ecoregion scale to align with Bird Conservation Regions which are delineated to include similar bird communities within each ecoregion (*64*). Our findings indicate that while communities within ecoregions experienced a gradual drift in species’ abundances, species richness remained stable. However, this broad scale may overlook context-dependent mechanisms occurring at more local scales (e.g., local abundance heterogeneity observed in (*48*)). Regarding network dynamics, a finer resolution may capture potential spatial heterogeneity within ecoregions but it may also exacerbate the impact of sampling on the representativeness of communities (*65*). Investigating network change at regional to continental scales is crucial as avian biodiversity patterns, trends, and drivers can differ markedly across spatial resolutions (*5*, *6*, *48*, *57*).

## Conclusion

Here, we show that North American bird assemblages have dramatically changed; a bird community in the 1970s differs from one observed in the 2020s in both species composition and network functions. These changes in bird community structure and function are correlated with impacts of human activity at regional to continental scales, providing evidence for the long-term and widespread extent of human presence impacts on wildlife (*59*). Our approach, which incorporates a metanetwork, network analysis, and spatiotemporal observations, is fully transferable to other taxonomic groups, providing a framework for other species and regions. By integrating comprehensive knowledge of interspecific bird interactions, we demonstrate that species interactions are important facets of biodiversity that can drive the functional organization of species within networks at regional to continental scales, and thus provide important information for monitoring and conservation prioritization.

## Acknowledgments

We dedicate this research to the memory of our young colleague, Dr. Marc Ohlmann, whose insights about ecological networks inspired this work. Marc initiated this new generation of ecological researchers who are as interested in ecology as they are in mathematics and philosophy. Marc had only just begun working on this project when the mountain took him away. This research would not be possible without the time and effort by numerous scientists, students, and volunteers that compiled the data. Specifically, we thank the Cornell Lab of Ornithology, researchers, and students involved in compiling the AvianMetaNetwork for North America. We also thank the many North American Breeding Bird Survey (BBS) volunteers for collecting data and USGS BBS scientists Jeff Hostetler, David Ziolkowski, and Jim Hines for helpful insights. We thank Lala Kounta for assistance with the ERA5 and Human Modification data, and Pat Bills, Kelly Kapsar, Lucas Mansfield, and Jenna Baljunas for enhancing the AvianMetaNetwork data workflow.

## Funding

French CNRS (VM, SD)

Michigan State University Ecology Evolution, and Behavior Seed Grant (PLZ)

## Author contributions

Conceptualization: PLZ, SD, VM

Methodology: VM, SD, PLZ

Investigation: PLZ, SD, VM

Visualization: SD, VM

Software: VM, SD

Data Curation: PLZ

Funding acquisition: PLZ

Project administration: PLZ

Writing – original draft: PLZ, SD, VM

Writing – review & editing: PLZ, SD, VM

## Competing interests

Authors declare that they have no competing interests.

## Data, code, and materials availability

The Human Modification data (*58*), ERA5 data (*66*, *67*), BBS data (*54*), and BCR data (*64*) are all publicly available. R version 4.5.2 was used to run analyses. The data from AvianMetaNetwork for North America (*43*) are available for reviewers at this EDI repository (*37*): P. L. Zarnetske, P. Bills, K. Kapsar, L. Mansfield, E. Parker, The AvianMetaNetwork: biotic interactions among birds of the continental United States and Canada, Environmental Data Initiative (2026) ver 2; https://doi.org/10.6073/pasta/9bc99f67618359b2d9a6770eff22664a.

All code and data to run the analysis and reproduce figures in this study is available at the following Environmental Data Initiative (EDI) repository (*84*): P. L. Zarnetske, S. Dray, V. Miele, Spatiotemporal network analysis of the AvianMetaNetwork across North America, 1970-2023, Environmental Data Initiative (2026) ver 2; https://doi.org/10.6073/pasta/03b472c365493cc92dc0a6fa704cc5bf. In the repository we include two QMD files and their rendered PDFs that reproduce the R analysis and figures, including supporting information figures, plus additional BCR-level result figures.

Both EDI repositories linked above are embargoed until publication.

## Supplementary Materials

### Materials and Methods

#### Study area and taxa

We focus on 687 breeding birds of the continental United States and Canada. Since 1966, breeding season point counts have been conducted through the North American Breeding Bird Survey (BBS), providing extensive spatial and temporal observations of birds across the continent (*54*). These breeding bird species are composed of 24 orders, 81 families, and 325 genera (*54*), and provide numerous ecosystem functions and services including seed dispersal, predation of pests, contributions to nutrient cycling within and across ecosystems, food for other fauna including humans via hunting, and recreation and human well-being via birding, nature walks, and ecotourism (*32*). The 687 species considered within this study result from a combination of the focal species within AvianMetaNetwork and observed BBS species, as detailed below.

#### Avian MetaNetwork for North America

The AvianMetaNetwork for North America is a digital natural history dataset consisting of 25,907 bird-bird interactions, involving 731 focal avian taxa occurring in the continental United States (including Alaska) and Canada plus 1,258 additional avian taxa interacting with the focal species during migration or in more southern portions of their ranges (*37*). These ‘focal taxa’ are determined from the combination of checklists (North American Breeding Bird Survey 2024 species list (*54*) and Avibase v8.17 regional checklists for Alaska, Canada, and the contiguous United States (CONUS) (*69–72*)), with taxonomy harmonized to the 2024 Clements/eBird checklist (*73*). These bird-bird interaction data result from manual systematic interpretations of interactions from text within species accounts of Birds of the World (BOW)(*33*)—the most comprehensive and curated set of avian species accounts. The reproducible workflow to build this database is outlined at (*37*, *43*) and follows the Environmental Data Initiative (EDI) data harmonization scheme for Levels 0 (raw), 1 (cleaned), and 2 (analyzed) data (*74*). Variables relevant to this study are described below.

Avian interactions inferred from text descriptions in BOW include competition over food and space, predator-prey interactions, brood parasitism, commensalism, mixed flocking, kleptoparasitism, and more. The 23 sub-types of interactions fall into seven main types with combinations of 0,1,-1 effects on each taxa in a pair: trophic, competition, facilitation, commensalism, parasitism, mobbing, and amensalism; other mainly neutral (0,0) interactions also occur (table S8). Each row in the interaction data represents a unique bird–bird interaction from a unique text excerpt within BOW, with attributes including: scientific and common names for each pair of avian taxa, interaction type, effect of the interaction on each species (negative (−1), positive (1), or neutral (0)), season of the interaction (non-breeding or breeding), the text excerpt source, and the exact text excerpts providing the interaction context.

In this study, we used species-level interaction data and observations during the breeding season to ensure that all interactions involved the same taxonomic level and timeframe. We subset the interaction data to align with the scope of the BBS (*54*); we omitted interactions that occurred during the non-breeding season, and only used interactions involving species observed in the BBS. We removed: 18 Genus-level taxa (e.g., *Buteo* sp.); 44 uncertain taxa designated with “/” where the species could be one or another species (e.g., *Aphelocoma californica/woodhouseii*); and 7 hybrid taxa that were rarely observed in the BBS or in the interaction data (e.g., *Anas platyrhynchos* x *rubripes*). We then grouped any subspecies into species, given that interactions were rarely found in the literature at the subspecies level and subspecies were rarely in the BBS observations. As AvianMetaNetwork includes multiple instances of an interaction sub-type between two species because it was documented in multiple text-based sources, we omitted duplicate pairwise interactions among a given species’ pair to keep only unique pairwise interactions per species pair.

We created six main ecological networks by aggregating all unique sub-types of interactions to their interaction type (see table S8). We removed amensalism due to the rarity of this interaction type (50 records). We also did not include neutral species relationships for which there is no known direction or effect (0,0; 2728 records). These neutral relationships include courtship, hybridization, play, and co-occurrence, and are not considered species interactions per se. Finally, we finally removed the few recorded cases of intra-specific interactions (6 observations).

The final data subset of unique pairwise interactions among species for use in network modeling was 13,762 interactions among 687 focal North American species.

#### Network-based functional groups

We used the 13,762 interactions among all 687 focal North American species to compute networks containing potential interactions among species in the study area. We considered the six networks independently for the six different types of interactions. For each network, we defined the “non-interacting” group as the subset of the 687 species that do not appear in the interaction pairs, i.e., that are not active for the type of interaction. After removing this group for a given network, we used network clustering (*50*) to gather species in functional groups, i.e., groups of species that tend to behave in the same way in terms of interaction involvement and interaction partners. We used stochastic block models (SBMs;(*75*)), already used in ecology for different types of interactions (e.g., (*39*)). With a SBM, species (nodes of a graph) are assigned to homogenous sets using interaction probabilities, without prior knowledge. The number of groups is jointly selected, with the Elbow method which consists of detecting the point where the likelihood slope decreases to reach a plateau. Using the blockModels R package v1.1.5 (*76*), we ultimately obtained six network-based functional groups of species, each associated with one of the six networks.

#### Bird observations and ecoregions

We used BBS count observations from 1970 to 2023 (*54*), excluding 1966-1969 when there was low sampling, and 2020 due to no data collection during the COVID-19 pandemic. The BBS was established to monitor annual bird population trends during the breeding season across North America, using a standardized protocol across established routes. BBS data consist of routes that are surveyed once yearly at the peak of the breeding season (April-June); around 500 routes were surveyed in the beginning of the survey, and in more recent decades, around 3000 routes are surveyed every year. During a survey, a qualified amateur observer conducts a 3-minute point count every 0.5 miles (∼800 m) along the 24.5-mile (∼40 km) route (50 stops per route), recording the number of individuals seen or heard of each bird species within a 400m radius. Although routes have been added over time, and some routes have been retired, the majority of routes have been consistent throughout the survey. In our ecoregion-scale analysis, we use BBS count data as a proxy for abundance.

To reduce bias in our analysis, we only used BBS route-year combinations where the observer followed the official BBS protocol (as indicated by RunProtocolID = 101), resulting in a subset of ∼ 98% of the BBS data over the entire timeframe. We then selected species for which there are more than 50 BBS observations in total over the entire time period 1970-2023 (*54*). To reduce any effect of unequal sampling among ecoregions and across years because of the number BBS routes can vary over time and space (*54*, *77*), we divided the raw numbers of observations across all species, by the number of routes for every year and every ecoregion.

To evaluate different bird communities spatially across different ecoregions of North America, we used ecoregion boundaries defined by the Bird Conservation Regions (BCRs) (*64*). We excluded BCR regions with <10 routes for all or part of the period; these regions are in the northernmost portion of North America and lack the higher geographic density and frequency of BBS route visits that are more typical of the lower 48 United States. The 5 excluded BCRs include: 1 (Aleutian/Bering Sea Islands); 2 (Western Alaska); 3 (Arctic Plains and Mountains); 4 (Northwestern Interior Forest); and 7 (Taiga Shield and Hudson Plains).

The final observation data subset for this analysis included 541 species and 5,431 routes, for a total of 98,003,061 route-level observations over 1970-2023, across 32 ecoregions.

#### Analysis of temporal community patterns

##### Species’ population-level

We considered an additional normalization to remove the effects of difference in total abundances across species and thus focus simultaneously on temporal variations of rare to ubiquitous species. We rescaled abundances between 0 and 1 with a min-max transformation for each region, species by species. A value of 1 indicates an observation peak for a particular species (maximum over the entire time period in this region), and a 0 indicates an observation minimum, respectively. With this normalization, all species have the same weight in the remaining analysis, ensuring that highly abundant species do not dominate the results. We obtained a series of 32 tables (one per BCR) containing the min-max transformed abundances of species (columns) for each year (rows).

##### Network-level and Network Group-level

We worked with the overall number of observations summed per network group, including the groups of non-interacting species, for each type of interaction independently. In other words, we built a community vector for which the cells are the abundance of each network group. However, we also normalized the abundance per year such that they sum to 1, in order to compare group proportions instead of group row abundances that can be driven by over-abundant species. With a similar reasoning than at the species level, we performed a min-max transform per group to consider the network groups on equal footing. We obtained a series of 32 tables (one per BCR) containing the min-max transformed abundances of the network groups (columns) for each year (rows).

##### Multivariate analysis with MFA

We analyzed both series of tables (at species’ population and network levels) similarly, using Multiple Factor Analysis (MFA; (*78*)) with the ade4 R package v1.7-23 (*79*). At the species level, this multivariate method first identified the reference structure which best summarizes the main trends shared by species across all BCR. This step consisted of performing a PCA of the super-table concatenating the series of sub-tables (one per BCR) stacked horizontally. In this PCA, each sub-table was down-weighted according to the first eigenvalue of its PCA so that BCRs with more species or more variability do not dominate the global analysis. As with PCA, MFA produces global components and scores for years and species per BCR so that it is possible to identify the main structures in the data. It is important to note that the trends resulting from the analysis are not necessarily related to the timeline: years are compared independently to their order and any chronological ordering is solely a result of the analysis. For the first MFA global component, we reported the (i) scores for years to characterize global temporal trends, (ii) the average scores per species to identify species that are involved (or not) in these temporal changes and (iii) the links (that varies between 0 and 1 and measure the amount of inertia of each BCR sub-table projected on the global component) to study the spatial variations of temporal structures and evaluate how each BCR follows or not the global temporal trends. In other words, the link value for a given BCR measures the agreement between the trend at the global scale and what is actually happening in this BCR. At the network level, the analysis is the same except that species are replaced by network groups.

##### Climate and human modification analysis

To investigate the potential influence of changes in climate and human impacts on species-level and network-level turnover in different BCRs across the timeframe, we quantified the Pearson correlation coefficient (and evaluated the significance of the associated t-tests by providing critical values) between each turnover and climate variables, and between each turnover and human impact variables, on a per-BCR basis. WGS84 (latitude, longitude) was used as the geographic coordinate reference system for all maps and spatial analyses.

To document changes in climate, we used two climate variables that are known to influence the composition of ecological communities including birds: temperature and precipitation; (*80–82*). Specifically, we used the yearly mean annual surface temperature (degrees C at 2m height above ground) and the yearly total precipitation (mm/year) for each year (1969–2023) from the ERA5 climate reanalysis data (0.25×0.25 degree gridded data; (*66*, *67*)). We defined “year” as July-June to align with the BBS year data (collected May-early July (*54*)), such that the months leading up to the BBS observations were mostly included in that observation year, whereas the months following were largely included in the subsequent BBS observation year. Per grid cell, we then computed linear regressions for temperature, and for precipitation, over the 1970-2023 period. The slopes of these regressions per grid cell were then used to obtain BCR-level mean slope values by taking the area-weighted average of the grid slopes across each BCR (applying area weighting to account for unequal grid cell sizes). On a per-BCR basis, we correlated the slopes of the temperature and precipitation change with each of the turnovers (species-level and network group-level), resulting in continental-scale Pearson correlations between changes in climate variables and each turnover.

For each BCR, we used temporal change in the Human Modification data (300m resolution gridded data; (*58*, *68*)) to assess the correlation between threats facing species and turnover. These threat data incorporated into the modification data follow the structure of the IUCN threat framework (*83*), resulting in 8 modification variables: Residential, commercial, and recreation areas (BU; i.e., footprint associated with built-up land uses), Agriculture (AG), Energy production & mining (EX), Transportation & service corridors (TI; i.e., road and railways, power lines, nighttime lights), Biological resource use (FR; i.e., footprint of timber harvesting), Human accessibility (HA; human intrusion i.e., use of roads or trails backcountry recreation, or risks for invasive species), Natural systems modification (NS; i.e., dams and reservoirs) and Pollution (PO). The timeframe available for these data is 1990-2020, reflecting a large portion of our study. During this timeframe, data are available as five-year change measures of each of the modification variables (1990, 1995, 2000, 2005, 2010, 2015, and 2020). Per 300m grid cell, a value of 0 indicates no modification change over the prior 5 years, and 1 is complete modification change over the previous 5 years. Per grid cell, we summed the modification values per variable across all 5-year periods and obtained a global modification change value for each threat. Per BCR, we then took the area-weighted mean of the grid-level summed modification change across all grid cells within a BCR (with area-weighting applied to grid cells to account for unequal grid cell sizes across the study area). On a per-BCR basis, we then correlated each of the 8 modification change values with each of the turnovers, resulting in continental-scale Pearson correlations between cumulative changes in human modification and each turnover.

**Fig. S1.**
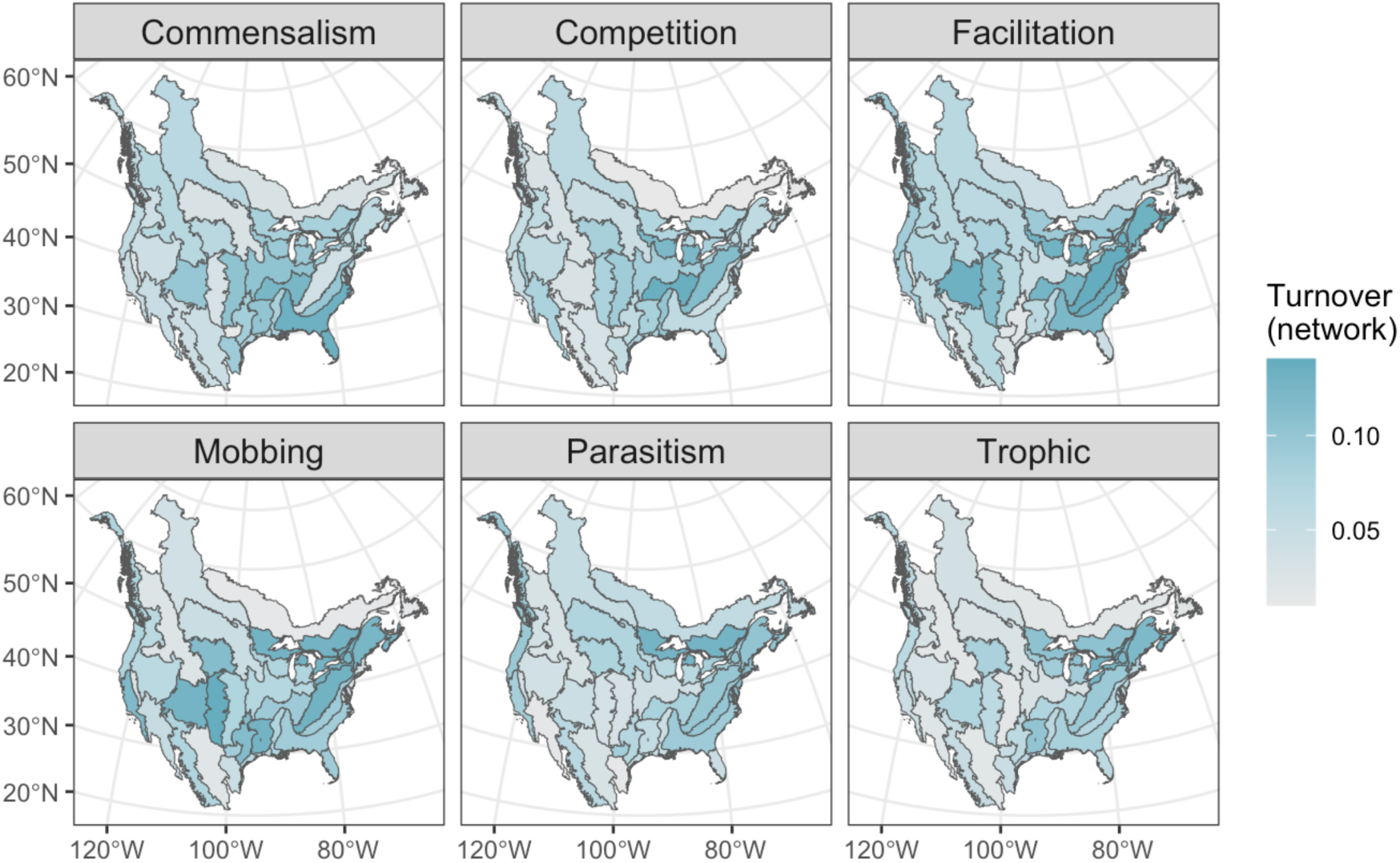
Network turnover by interaction type and Bird Conservation Region (BCR) (*64*). Network turnover was computed for each BCR and interaction type by decomposing the global measure of the temporal turnover (computed from the multifactor analysis (MFA) coordinates) to identify the variation partition per interaction type and BCR. The breakdown of the first MFA eigenvalue into percentages by type of interaction (reflecting each network’s overall contribution to turnover) is: mobbing (20.64%), commensalism (18.16%), competition (17.10%), parasitism (16.29%), trophic (15.29%), and facilitation (12.53%).

**Fig. S2.**
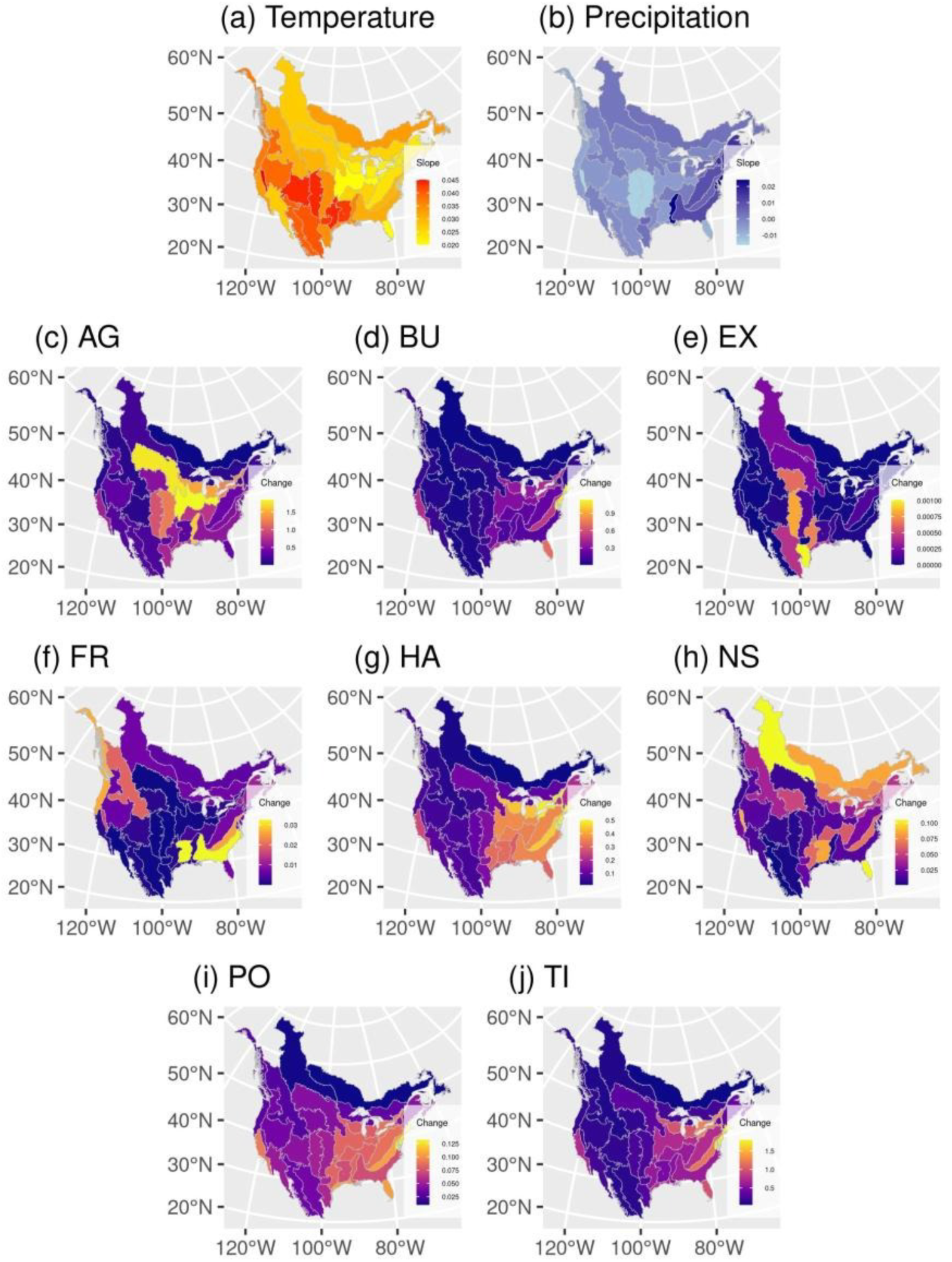
Climate and human modification changes by 32 Bird Conservation Regions (BCR) (*64*). Changes in climate from 1969-2023 are shown by (a) the slope of mean annual surface temperature (degrees C at 2m height above ground), and (b) the slope of total annual precipitation (mm/year) (from (*67*)). Human modification (HM) change is shown for 8 different variables as the BCR area-weighted mean of the cumulative modification change from 1990-2020 (from (*68*)). Per 300m human modification grid cell, a value of 0 indicates no modification change over the prior 5 years, and 1 is complete modification change over the previous 5 years. The 8 HM variables are: (c) AG: agriculture; (d) BU: residential, commercial, and recreation areas, or the footprint associated with built-up land uses; (e) EX: energy production & mining; (f) FR: biological resource use (i.e., footprint of timber harvesting); (g) HA: human accessibility and intrusion, i.e., use of roads or trails backcountry recreation, or risks for invasive species; (h) NS: natural systems modification (i.e., dams and reservoirs); (i) PO: pollution; and (j) TI: transportation & service corridors, i.e., road and railways, power lines, nighttime lights. From 1970 to 2023, mean annual temperature increased in all ecoregions, and total precipitation increased in some ecoregions but decreased in others (a-b). Over this time, the US Southwest experienced the highest increase in temperature and some of the largest decreases in total precipitation, whereas many of the Eastern regions experienced the largest increases in total precipitation and moderate temperature rise. Regarding human modification data in the period 1990-2020 (c-j), the overall human activity changes (e.g.; HA; TI, PO, BU) are more pronounced in the Eastern regions. On the other hand, agricultural and exploitation changes are more noticeable in some Central regions. WGS84 (latitude, longitude) was used as the geographic coordinate reference system for all maps and spatial analyses. See Materials and Methods for more details.

**Fig. S3.**
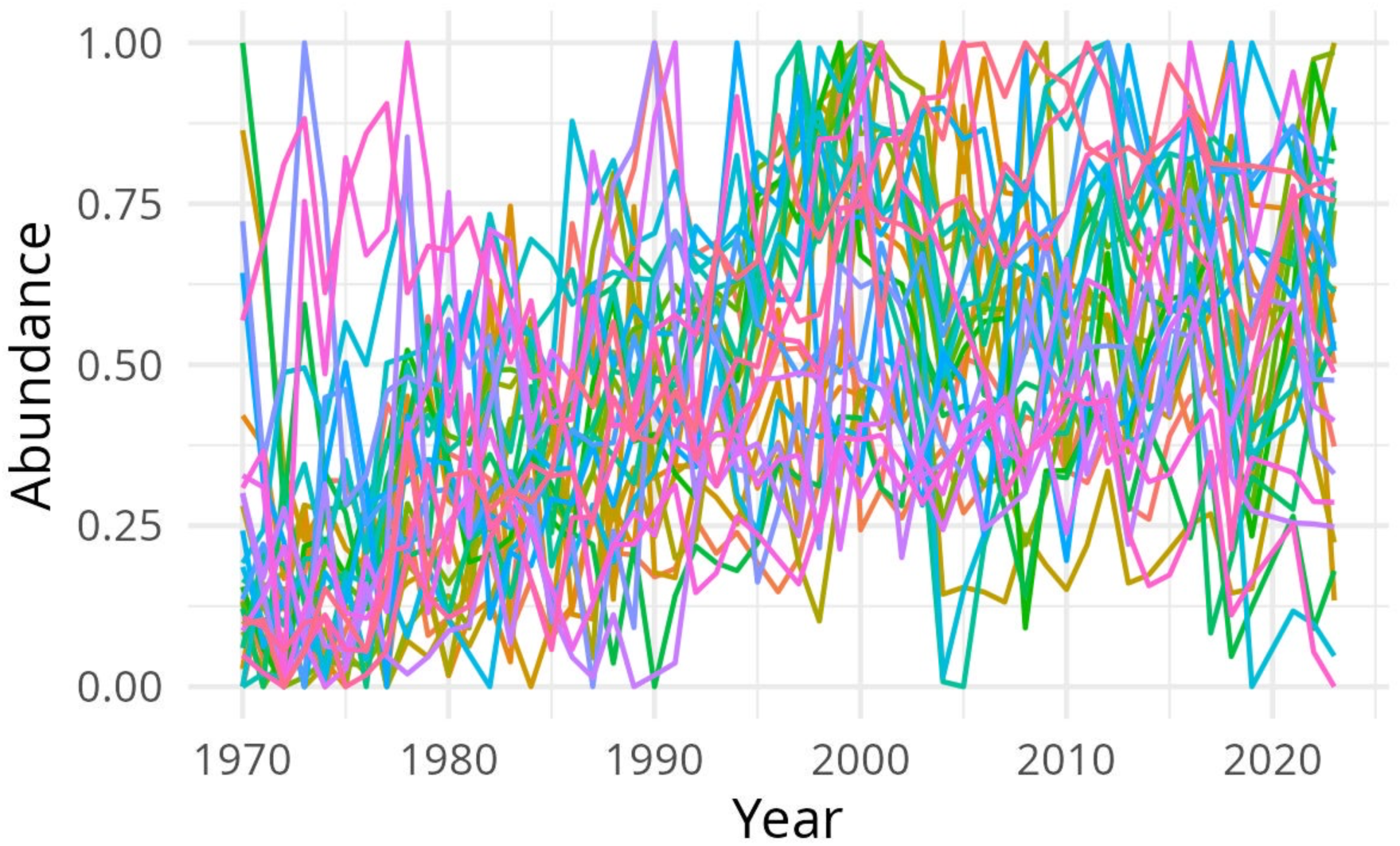
Abundance trends for the bird species (each color is a species) in the top predator group (orange nodes in the trophic network in Fig. 3).

**Table S1.** Interaction probabilities (percents) between the Trophic groups obtained with Stochastic Block Models for each interaction type. See Fig. 3 legend for descriptions of group color membership of different species.

| **Trophic** | green | orange | white |
| --- | --- | --- | --- |
| green | 2.8 | 2.0 | 3.3 |
| orange | 19.6 | 27.5 | 15.4 |
| white | 0.0 | 0.0 | 0.1 |

**Table S2.** Interaction probabilities (percents) between the Commensalism groups obtained with Stochastic Block Models for each interaction type. See Fig. 3 legend for descriptions of group color membership of different species.

| **Commensalism** | green | orange | purple | white |
| --- | --- | --- | --- | --- |
| green | 1.3 | 0.7 | 1.4 | 0.0 |
| orange | 9.0 | 3.8 | 4.4 | 0.6 |
| purple | 0.0 | 0.3 | 0.8 | 0.0 |
| white | 0.0 | 0.1 | 2.0 | 0.3 |

**Table S3.** Interaction probabilities (percents) between the Parasitism groups obtained with Stochastic Block Models for each interaction type. See Fig. 3 legend for descriptions of group color membership of different species.

| **Parasitism** | green | orange | purple | white |
| --- | --- | --- | --- | --- |
| green | 6.8 | 3.4 | 0.0 | 0.8 |
| orange | 44.3 | 0.0 | 3.0 | 41.0 |
| purple | 0.2 | 0.0 | 2.2 | 0.1 |
| white | 0.2 | 0.0 | 0.0 | 0.1 |

**Table S4.** Interaction probabilities (percents) between the Competition groups obtained with Stochastic Block Models for each interaction type. See Fig. 3 legend for descriptions of group color membership of different species.

| **Competition** | green | orange | purple | white |
| --- | --- | --- | --- | --- |
| green | 16.8 | 0.5 | 0.2 | 0.0 |
| orange | 0.5 | 11.8 | 0.1 | 2.0 |
| purple | 0.2 | 0.1 | 1.8 | 0.1 |
| white | 0.0 | 2.0 | 0.1 | 1.0 |

**Table S5.**
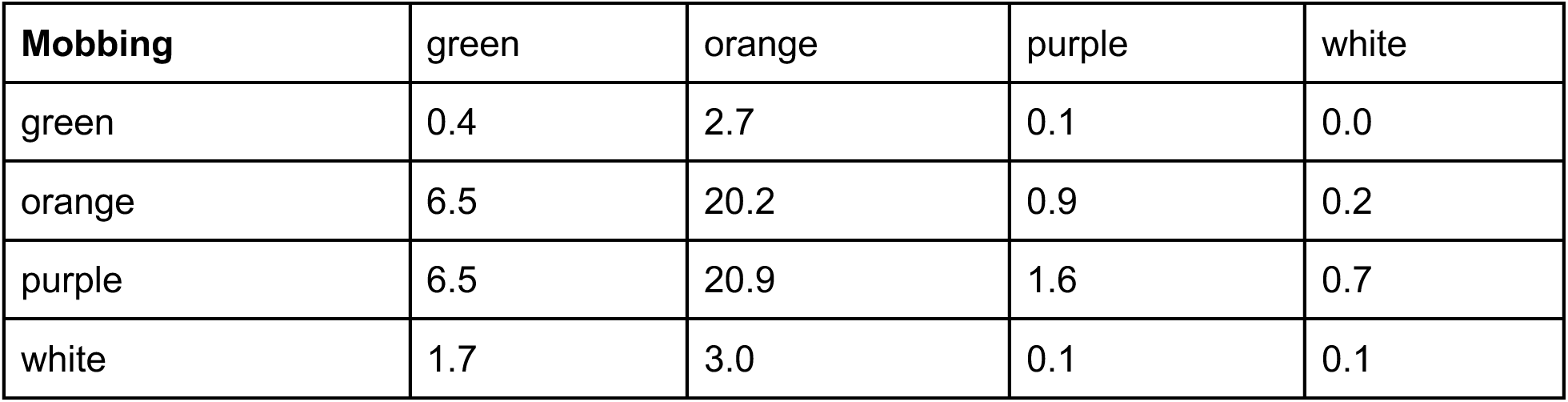
Interaction probabilities (percents) between the Mobbing groups obtained with Stochastic Block Models for each interaction type. See Fig. 3 legend for descriptions of group color membership of different species.

| **Mobbing** | green | orange | purple | white |
| --- | --- | --- | --- | --- |
| green | 0.4 | 2.7 | 0.1 | 0.0 |
| orange | 6.5 | 20.2 | 0.9 | 0.2 |
| purple | 6.5 | 20.9 | 1.6 | 0.7 |
| white | 1.7 | 3.0 | 0.1 | 0.1 |

**Table S6.** Interaction probabilities (percents) between the Facilitation groups obtained with Stochastic Block Models for each interaction type. See Fig. 3 legend for descriptions of group color membership of different species.

| **Facilitation** | green | orange | purple | white |
| --- | --- | --- | --- | --- |
| green | 100 | 0.2 | 4.5 | 0.2 |
| orange | 0.2 | 64.6 | 3.0 | 0.3 |
| purple | 4.5 | 3.0 | 7.0 | 0.1 |
| white | 0.2 | 0.3 | 0.1 | 1.3 |

**Table S7.** Interaction networks and their corresponding interaction effects, types, and definitions. In this study we do not include amensalism due to its rarity among species. We do not include interactions or relationships between species where the interaction has an unknown effect on one or both of the interacting species or are not considered species interactions in a community ecology context per se (Interaction Type “Other”: co-occurrence, play, hybridization, copulation, courtship, accidental killing).

| **Interaction Type** | **Interaction Effects** | **Interaction Sub-Type** | **Interaction Definition** |
| --- | --- | --- | --- |
| Trophic | -1,1 | Predation | Predator eats prey; predator = 1, prey = -1. |
| Competition | -1,-1 | Competition | Competing over space, territory, food, etc.; both species = -1. |
| Facilitation | 1,1 | Facilitation | When both interacting species benefit (if there is no specific facilitation interaction), e.g., creching, mutual allopreening; both species = 1. |
| Facilitation | 1,1 | Facilitation-mixed flocking | Special case of facilitation when both species benefit by being in a group; both species = 1 |
| Facilitation | 1,1 | Facilitation-comigration | Special case of facilitation when species migrate together; both species = 1 |
| Facilitation | 1,1 | Facilitation-feeding | Special case of facilitation when both species feed together, either in mixed-flocks or feeding aggregations; both species = 1 |
| Facilitation | 1,1 | Communal nesting | Special case of facilitation when species nest together, usually during breeding season; both species = 1. |
| Facilitation | 1,1 | Communal roosting | Special case of facilitation when species roost together, typically during nonbreeding season; both species = 1. |
| Commensalism | 0,1 | Commensalism | One interacting species benefits and the other receives no effect (example: cavities are constructed by primary cavity nesting bird like a woodpecker, and later used by another bird for its own nest (without any conflict)); species that benefits from the interaction = 1; species that provides the benefit = 0. |
| Commensalism | 0,1 | Commensalism-scavenge | Special case of commensalism when a species consumes/scavenges a previously deceased species or deserted/addled eggs; scavenger=1, dead species=0. |
| Commensalism | 0,1 | Commensalism-chick adoption | Special case of commensalism when one species adopts another species' chicks/eggs; species that is adopting = 0, species that is adopted = 1. |
| Commensalism | 0,1 | Commensalism-call mimicry | Special case of commensalism when one species copies the call/song of another; species that is mimicking = 1, species that is being mimicked = 0. |
| Parasitism | 1,-1 | Kleptoparasitism | Special case of parasitism when one species steals food or nest material from another species; species that steals and benefits = 1, species that is stolen from = -1. |
| Parasitism | 1,-1 | Brood parasitism | Special case of parasitism when one species places its egg in another species' nest so the nesting species has to raise a conspecific (nest parasitism - includes egg-dumping); species that lays egg in another species nest = 1, species that has to care for the conspecific young = -1. |
| Parasitism | 1,-1 | Nest takeover | Special case of parasitism when one species takes over (steals) the nest of another species, sometimes kicking out the existing eggs; species that takes over = 1; species whose nest is stolen = -1. |
| Amensalism | 0,-1 | Amensalism | One species causes harm to another species without any cost or benefits to itself (e.g. flushing from nest in response to perceived predator, accidental destruction of eggs); harmful species = -1, harmed species = 0. |
| Mobbing | 0,-1 | Mobbing | Usually indicative of predation, where the smaller species mobs the bigger species; the species doing the mobbing (usually potential prey) = 0; the species receiving the mobbing = -1 (the potential predator). The interaction is coded this way because mobbing is almost always deterring a predator, which reduces the likelihood of predation. |
| Other | 0,0 | Co-occurrence | Not an interaction per se. Species occur together but it is not clear what the interaction is; each species = 0. |
| Other | 0,0 | Play | Species play with each other; both species = 0. |
| Other | 0,0 | Hybridization | When species successfully mate and produce hybrid offspring; includes intergrades (hybrids of subspecies); both species = 0. |
| Other | 0,0 | Copulation | Unsuccessful mating attempt between two individuals of different species; both species = 0. |
| Other | 0,0 | Courtship | One species attempts to court another; both species = 0 |
| Other | 0,-1 | Accidental Killing | when one species accidentally kills another, but has no intent on doing so (e.g., no intent on preying on the dead species or competing with the dead species); species killed = -1, species who causes the death = 0 |

## Notes

https://doi.org/10.6073/pasta/2a13796a5fe8da1e650f08abc53e013f

### Competing Interest Statement

The authors have declared no competing interest.

### Summary of Updates

This version of the manuscript is more concise and has been reworked from the previous version.

